# Reduced mesophyll conductance by cell wall thickening and chloroplasts decreasing driven the decline of photosynthesis under sole NH_4_^+^ supply

**DOI:** 10.1101/2022.01.05.475041

**Authors:** Yiwen Cao, Yonghui Pan, Tianheng Liu, Min Wang, Shiwei Guo

**Affiliations:** Jiangsu Provincial Key Lab for Organic Solid Waste Utilization; National Engineering Research Center for Organic-based Fertilizers; Jiangsu Collaborative Innovation Center for Solid Organic Waste Resource Utilization; Nanjing Agricultural University, Weigang 1, Nanjing 210095, China

**Keywords:** Ammonium, leaf anatomy, *Lonicera Japonica*, mesophyll conductance, N form, nitrate, photosynthetic limitations, shrub

## Abstract

The relationship between nitrogen (N) sources and photosynthetic capacity of leaf differs between species. However, the leaf anatomical variabilities related to photosynthesis (*A*) of shrubs under different forms of N remain imperfectly known. Here, *Lonicera Japonica* (a shrub) was grown hydroponically in the presence of three forms of N (sole NH_4_^+^, 50%/50% NH_4_^+^/NO_3_^−^ and sole NO_3_^−^). *A* and photosynthetic N use efficiency significantly decreased under sole NH_4_^+^ supply, in parallel with down-regulated stomatal conductance (*g*_s_), mesophyll conductance (*g*_m_), and electron transfer rate (*J*). Up to the total *A* decline of 41.28% in sole NH_4_^+^ supply (compare with sole NO_3_^−^), the *g*_m_ attributed to 60.3% of the total limitations. Besides, the decreased internal air space explained the increase of gas-phase resistance, and the increased liquid-phase resistance in sole NH_4_^+^ supply was ascribed to the thicker cell wall thickness (*T*_cw_) and decreased chloroplasts exposed surface area per unit leaf area (*S*_c_/*S*). The discrepancy of *S*_c_/*S* could be interpreted by the altered chloroplasts numbers and the distance between adjacent chloroplasts (*D*_chl-chl_). These results indicate the alteration of *T*_cw_ and chloroplast numbers were the main causes of the difference in *g*_m_ in coping with varied N sources.

**Highlight:** Cell wall and chloroplast variability determining the mesophyll conductance under different nitrogen forms

## Introduction

*Lonicera Japonica* is a widespread semi-deciduous shrub species of the Caprifoliaceae family utilized in traditional medical practices (Schierenbeck, 2004; Shang *et al*., 2011). The stomatal and photosynthetic function are suggested the key traits for enhancing yield and quality of medicinal plants, as the photosynthesis improvement could increase carbon fixation and the production of biomaterials. Mesophyll conductance (*g*_m_) represents the diffusion conductance to CO_2_ in the mesophyll tissues from the sub-stomatal cavities to the carboxylation sites inside chloroplast (Peguero-Pina *et al*., 2012), which was proposed to be the first limiting factor in photosynthesis (Lu *et al*., 2016; Ren *et al*., 2019).

*g*_m_ is determined by the CO_2_ diffusion characteristic. CO_2_ diffusion, according to Fick’s law, depends on CO_2_ diffusivity, leaf temperature, diffusion distance, and the nature of the media in which diffusion occurs (e.g., mesophyll tissues) (Flexas *et al*.,2012), thus it is not strange that even tiny leaf anatomical traits could drive *g*_m_ variability. Tholen *et al*. (2012a) had reported that altered leaf anatomy significantly influences the CO_2_ diffusion in leaves. Leaf mass per area (*M*_A_) is an important integrated leaf morphological characteristic that regulates photosynthesis by influencing *g*_m_ (Niinemets, 2001; Poorter *et al*., 2009). Early studies reported a negative correlation between *M*_A_ and *g*_m_ (Flexas *et al*., 2008; Niinemets *et al*., 2009b), and further in species with high *M*_A_, photosynthesis is more limited by *g*_m_, on average by 57% (Tomás *et al*., 2013); analyses on the relationship between *g*_m_ and leaf density/leaf thickness, two components of *M*_A_ also corroborated this idea, despite the influences are opposite. The increase of *M*_A_ due to increased leaf density might result in a densely packed mesophyll cell, which will ultimately reduce the *g*_m_ by decreased mesophyll surface area exposed to intercellular airspace per unit leaf area (*S*_mes_/*S*) (Flexas *et al*., 2008; M. Weraduwage *et al*., 2016).

In the journey of CO_2_ diffusion, the mesophyll resistance is comprised of gas-phase resistance (i.e., from the intercellular air space) and liquid-phase resistance (i.e., from the cell wall, lipid membrane, cytoplasm, stroma, and chloroplast envelope). In particular, liquid-phase limitation accounts for more than 80% of the total limitation, in which the cell wall thickness (*T*_cw_) and chloroplast surface area exposed to intercellular airspace per unit leaf area (*S*_c_/*S*) are the two dominating components that affect *g*_m_ (Flexas *et al*., 2021). There are ample studies showing a significant positive correlation between *S*_c_/*S* and *g*_m_, and the *S*_c_/*S* was considered the uppermost parameter in determining *g*_m_ (Hu *et al*., 2020; Tomás *et al*., 2013). While the cell wall is often negligible, because it accounts for a tiny fraction of the CO_2_ diffusion length (Carriquí *et al*., 2020), has a comparative larger pore size and variability (Carpita *et al*., 1979), and quite a static influence on photosynthesis (Evans *et al*., 2009). Interestingly, in *Lycopodiales* (a species of fern), the accountability of *T*_cw_ in robust leaves of the total *g*_m_ was up to 70% (Tosens *et al*., 2016). Other components, such as the mesophyll thickness, plasma membrane, and chloroplast density, despite not that important, also play an important role in regulating *g*_m_, as observed in different plant species (Lu *et al*., 2016; Niinemets *et al*., 2009b; Veromann-Jurgenson *et al*., 2020a; Veromann-Jurgenson *et al*., 2020b). Overall, the leaf anatomies might vary independently and compensate for each other to achieve substantial *g*_m_ in some cases (Peguero-Pina *et al*., 2017).

In fact, *g*_m_ is a rapidly adapting trait, and thus its value presents a large variability across species or environments. Up to now, mesophyll conductance to CO_2_ had been studied on various plant species in a scale of interspecific variation and diverse environmental conditions. The review of Flexas *et al*. (2008) summarized the *g*_m_ in different pooled groups of plants, which showed an apparent increasing tendency of *g*_m_ from conifers (slightly above 0.1 mol m^-2^ s^-1^) to grasses (0.2-0.4 mol m^-2^ s^-1^), and the semi-deciduous shrubs generally had a *g*_m_ value of 0.2-0.3 mol m^-2^ s^-1^. The varying environmental conditions, such as light, water availability, salinity, and temperature had also been studied in regulating *g*_m_ (Flexas *et al*., 2008; Niinemets *et al*., 2009b; Pons *et al*., 2009; Tosens *et al*., 2012a). In contrast, the effects of nutrition on *g*_m_, particularly on quantitative limitations of *g*_m_ are somewhat incomplete, and only recently have the effects of nutrition (e.g., potassium, nitrogen) been reported (Gao *et al*., 2020; Hu *et al*., 2020; Xie *et al*., 2020; Xiong *et al*., 2015b). Nitrogen (N) is an important nutrition element for plant growth and photosynthesis. There is sufficient evidence of the strong interplays between the photosynthetic process and plant endogenous N status (Perchlik and Tegeder, 2018; Xiong *et al*., 2015). N in soil is available mainly in two inorganic N sources, i.e., nitrate (NO_3_^−^-N) and ammonium (NH_4_^+^-N). Due to the different assimilation processes and energy consumption, or sometimes the toxic effect of NH_4_^+^, photosynthesis of species showed quite different responses/preferences to NH_4_^+^ or NO_3^-^_. In some studies, the plant supplied with NH_4_^+^ was observed increased chloroplast numbers and chloroplast volume in comparison with NO_3_^−^ (Golvano *et al*., 1982), yet the other study observed an elevated *S*_c_/Rubisco in NO_3_^−^ supply (Gao *et al*., 2020); this effect on photosynthetic capacity and *g*_m_, according to some studies, could be ascribed to N allocation trade-off and source-sink balance within leaves (Evans and Clarke, 2019; Hikosaka, 2004). The studies on the relative importance of N source to the response of *g*_m_ variation in a scale of leaf structural traits had been reported in herbs (Gao *et al*., 2020) and trees (Liu *et al*.,2021), while the study on shrubs is largely lost. Moreover, major limiting factors that restrict the leaf photosynthetic capacity of *L. Japonica* under different inorganic source is largely unknown.

In the present study, *L.Japonica* was grown in a hydroponic experiment with three forms of N (sole NH_4_^+^, 50%/50% NH_4_^+^/NO_3_^−^ and sole NO_3_^−^) to investigate: (1) the variations of leaf photosynthesis and leaf anatomy under different inorganic nitrogen sources; (2) the crucial role of *g*_m_ on leaf photosynthesis of *L. Japonica;* (3) the impact of mesophyll conductance on leaf photosynthesis through leaf structure variations.

## Material and Methods

### Plant material and experimental condition

The hydroponic experiment was conducted in a greenhouse with a light intensity of 1000 μmol photons m^-2^ s^-1^ at the leaf level using 14-h photoperiod and day/night temperature of 28/18□. The experimental site is located in Nanjing (118°51’E, 32°1’N), China. One-year-old seedlings of *Lonicera japonica* were transferred to a 12 L rectangular plastic box (64 cm×23 cm×18 cm) and a one-half-strength mixture of NH_4_^+^ and NO_3_^−^ nutrient solution was supplied (for composition, see below). After two weeks, the seedlings were supplied with full-strength nutrition solution for two weeks, after which the uniform seedlings were transplanted to a 12 L rectangular plastic box (64 cm×23 cm×18 cm). The nutrition solution contained 2.0 mM MgSO_4_·7H_2_O, 2.8 mM CaCl_2_·2H_2_O, 1.03 mM K_2_SO_4_, 0.32 mM KH_2_PO_4_, 9.10×10^-3^ mM MnCl_2_·4H_2_O, 0.52×10^-3^ mM (NH_4_)_6_Mo_7_O_24_·4H_2_O, 37×10^-3^ mM H_3_BO_3_, 0.15 ×10^-3^ mM ZnSO_4_·7H_2_O, 0.16 ×10^-3^ mM CuSO_4_·5H_2_O, 35.8×10^-3^ mM Fe-EDTA. In our preliminary experiment, plant growth was highest at a ratio of 50%/50% NH_4_^+^/NO_3_^−^ compared with 75%/25% NH_4_^+^/NO_3_^−^ and 25%/75% NH_4_^+^/NO_3_^−^ at a N level of 2.8 mM. Thus, the N was supplied at 2.8 mM with three different treatments: sole (NH_4_)_2_SO_4_ (A), sole Ca (NO_3_) _2_ (N) or mixed N (combination of 50% (NH_4_)_2_SO_4_ and 50% Ca (NO_3_)_2_, AN). CaCl_2_ was added to the solutions of A and AN treatment to adjust the Ca level to N treatment (2.8 mM). Dicyandiamide was added to each nutrition solution as nitrification inhibitor. The nutrition solution was aerated for 1-h/1-h day/night and renewed every 4 days, while the pH was adjusted to 6.0±0.1 each day. The containers were placed randomly to prevent the position effect.

### Gas exchange and chlorophyll fluorescence measurement

Leaf gas exchange and chlorophyll fluorescence were measured simultaneously using an open gas exchange system equipped with multiphase flash (LI-6800XT; LI-COR Inc., Lincoln, NE, USA) from 8:30 to15:00. For each treatment, new fully-expanded leaves were randomly selected for the measurements with five replications. Measurements were obtained at a leaf temperature of 28±0.5°C, CO_2_ concentration inside the chamber of 400±6 μmol mol^-1^, and a photosynthetic photon flux density (PPFD) of 1000 μmol photons m^-2^ s^-1^. For fluorescence parameters, steady-state fluorescence yield (*F*_s_) and maximum fluorescence (*F*_m_’) were recorded using a fluorometer chamber during a light-saturating pulse of approximately 8000 μmol m^-2^ s^-1^. During the photosynthetic CO_2_ responses (*A*/*C*_i_ curve) measurements, the ambient CO_2_ concentration (*C*_a_) was set in a series from 400 to 300, 200, 150, 100, and 50 μmol CO_2_ mol^-1^ and then increased from 50 to 400, 600, 800, and 1000 μmol CO_2_ mol^-1^ at a constant PPFD of 1000 μmol m^-2^ s^-1^ with four replicates. The maximum carboxylation rate (*V*_cmax_) was calculated according to Long and Bernacchi (2003), and the carboxylation efficiency (*CE*) was estimated as the slope of the *A/C*_i_ curve fitting line over a *C*_i_ range of 50-200 μmol mol^-1^. The effective quantum efficiency of photosystem II (ΦPSII) was quantified as follows: ΦPSII=(*F*_m_’-*F*_s_)/*F*_m_’. The potential electron transport rate (*J*) was calculated as *J*= ΦPSII×PPFD×α×β, where α is the leaf absorption and β is the proportion of quanta absorbed by PSII, assumed as 0.85 and 0.5, respectively. There were no differences in chlorophyll contents between A, N, and AN leaf, thus eliminating out the confounding effect of different leaf optical properties as a result of N forms among the treatments.

Then we estimated the mesophyll conductance (*g*_m_) and chloroplast CO_2_ concentration (*C*_c_) by variable *J* method proposed by Harley *et al*. (1992).

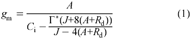

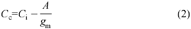

where *A*, *C*_i_, *C*_c_, and *J* were calculated as described previously, Γ^*^ is the CO_2_ compensation point in the absence of mitochondrial respiration and *R*_d_ is the mitochondrial respiration rate in the light. In this study, Γ^*^ was assumed to be 40.0 μmol m^-2^ s^-1^, and *R*_d_, according to the previous data, was assumed to be 1.0 μmol m^-2^ s^-1^.

### Leaf physiology

Four new fully expanded leaves from each treatment were randomly selected and pictured. The leaf area (*A*_L_) obtained from the picture was calculated by Image-Pro Plus (Media Cybernetics, Sliver Spring, MD, USA), after which the leaves were oven-dried at 105°C for 15 min and then dried to constant weight at 65°C and weighted. Dried leaf samples were weighed and digested with H_2_SO_4_-H_2_O_2_ at 270 °C, and the leaf N concentration was determined using the digital colorimeter (AutoAnalyzer 3; Bran+Luebbe). The total chlorophyll concentration was determined according to the method proposed by Sartory and Grobbelaar (1984). Approximately 0.5 g fresh leaf discs (avoiding the main vein) were extracted with 50 ml 95%(v/v) ethanol (analytically pure, Sinopharm Chemical Reagent Co., Ltd) in the dark until they were blanched (usually no more than two days). The extract solutions were used for the determination of chlorophyll *a* and *b* using a spectrophotometer (HBS-1096A, Shanghai, China) at 665 and 649 nm, respectively. The total chlorophyll was calculated as the sum of chlorophyll *a* and *b*. There were four replications for chlorophyll concentration determination.

### Quantitative limitation analysis of *A*

To separate the relative controls on *A* resulting from the difference in stomatal conductance, mesophyll diffusion, and biochemical capacity, quantitative analysis was conducted. The limitation of *A* under different types of N forms, according to Grass and Magnani (2005), was then composed of stomatal limitation (*S*_L_), mesophyll conductance (*MC*_L_), and biochemical limitation (*B*_L_), which could be expressed as follows:

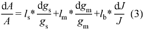

where the *l*_s_, *l*_m_, and *l*_b_ (*l*_s_+*l*_m_+*l*_b_=1) were the relative limitation of stomatal conductance, mesophyll diffusion and biochemical capacity, which were calculated as follows:

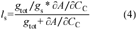

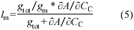

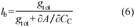

where the *g*_total_ is the total conductance to diffuse CO_2_ from the leaf surface (i.e., inside the leaf boundary layer) to the carboxylation site of chloroplast, which can be calculated as 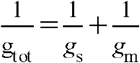. *∂A*/*∂C*_c_ was estimated as the slope of *A*/*C*_i_ curves over a *C*_i_ range of 50-100 μmol mol^-1^.

Then, the limitations of *A* can be approximately calculated as follows:

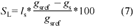

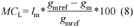

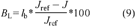

where *g*_sref_, *g*_mref_, and *J*_ref_ are the reference values of stomatal conductance (*g*_s_), mesophyll conductance (*g*_m_), and the potential electron transport rate (*J*), respectively. The reference values were obtained from the parameters from the optimal treatment from A, AN, and N. Here, we adopted the N treatment as the optimal treatment.

### Anatomical analysis

After the photosynthetic parameter measurements, leaf pieces (approximately 5mm×5 mm) were cut between the main veins from each of the four different plants for anatomical measurements and transmission electron microscope (TEM) analysis. For anatomical measurement, leaf segments were quickly fixed with FAA (95% ethanol: glacial acetic acid: formalin: distilled water =10:1:2:7, v/v) and dehydrated in dimethylbenzene-ethanol series. The pieces were then staining with safranin-fast green and embedded in paraffin. After cutting into 6 μm transverse sections, the pieces were photographed at a magnification of 400× with Nikon Eclipse E100 microscope equipped with a Nikon microscope camera (Nikon DS-U3). For TEM analysis, leaf materials were quickly fixed with glutaraldehyde (2.5%, v/v) in 0.1 M phosphate buffer (pH 7.4) under vacuum. Afterward, the segments were postfixed with 2% osmium tetroxide and dehydrated in a graded ethanol series, followed by washing in propylene oxide. The dehydrated segments were then embedded in Epon 812 resin. Ultrathin cross-sections were cut with a Power Tome-XL ultramicrotome, stained with 2% uranyl acetate, and then viewed under an H7650 transmission electron microscope (H-7650, Hitachi, Japan). Photos were taken at 2000-8000× direct magnification to measure the cell wall thickness and chloroplast characteristics. Leaf thickness (*T*_L_), leaf density (*D*_L_), leaf volume per area (*V*_A_=*A*_L_×*T*_L_), mesophyll thickness between two epidermises (*T*_mes_), and the volume fraction of intercellular air space (*f*_ias_) were calculated according to the light and TEM micrographs. The *D*_L_ and *f*_ias_ were determined as follows:

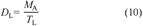

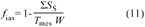

where *M*_A_ is the specific leaf weight (mg cm^-2^), Σ*S*_s_ is the total cross-section area of the mesophyll cells and *W* is the width of the measured framed range.

Chloroplast (*S_c_/S*) and mesophyll (*S*_mes_/*S*) surface area exposed to intercellular airspace per unit leaf area were also calculated from light and TEM micrographs, as reported by Evans *et al*. (1994) and Syvertsen *et al*. (1995).

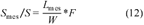

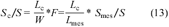

where the *L*_mes_ is the total mesophyll surface length facing intercellular air space per leaf area, and *L*_c_ is the chloroplast surface length facing intercellular air space per leaf area. *F* is the curvature correction factor, which depends on the shape of mesophyll cell and was calculated according to Thain (1983). Briefly, according to the differences of palisade and spongy cells (i.e., cell arrangement direction and axial/length ratio), *F* was calculated as a weight average of spongy (*F*_1_) and palisade (*F*_2_) mesophyll distributions:

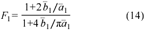

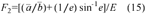

where 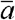 and 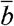 are the average of the length and thickness of the mesophyll. *e* is the eccentricity, and *E* is the elliptical integral, which were calculated as follows, respectively:

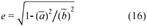

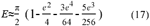

Besides, cell wall thickness (*T*_cw_), chloroplast length (*L*_chl_) and thickness (*T*_chl_), the distance between two neighboring chloroplasts (*D*_chl-chl_), chloroplast distance from the cell wall (*L*_cyt_), and the chloroplast number per mesophyll cell were measured from TEM micrographs at a 2000-8000×. The images were analyzed with Image-Pro Plus software (Media Cybernetics, Sliver Spring, MD, USA).

### *g*_m_ modelled from anatomical characteristics

To determine *g*_m_, the one-dimension gas diffusion model modified by Tomas *et al*. (2013) was applied in this study. *g*_m_, a composite conductance, is shared by different leaf anatomical characteristics and decided by within-leaf gas and liquid components:

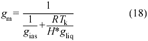

where *g*_ias_ is the gas-phase conductance which stands for the gas-phase pathway from substomatal cavities to the outer surface of cell wall, and *g*_liq_ is the liquid conductance that from the outer surface of cell wall to chloroplast. *R* is the gas constant (Pa m^3^ K^-1^ mol^-1^), *T*_k_ is the absolute temperature (K), and *H* is the Henry’s law constant (Pa m^3^ mol^-1^). Due to the *g*_m_ in this equation, is defined as a gas-phase conductance, thus *H*/*RT*_k_ (dimensionless form of Henry’s constant) is needed to convert to *g*_liq_ to the corresponding gas-phase equivalent conductance (Niinemets and Reichstein, 2003).

In this model, gas-phase conductance depends on gas-phase porosity (*f*_ias_) and effective diffusion path in the gas-phase (Δ*L*_ias_):

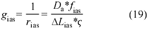

where ς is the diffusion path tortuosity (m m^-1^) and Da (m^2^ s^-1^) is the diffusion coefficient for CO_2_ in the gas-phase (1.51×10^−5^ at 25°C). Δ*L*_ias_ is taken as half of the mesophyll thickness (*T*_mes_).

The total liquid-phase diffusion conductance was determined by different components in mesophyll cell, including the conductance in the cell wall (*g*_cw_), plasma membrane (*g*_st_), cytosol (*g*_cyt_), chloroplast envelope (*g*_en_), and stroma (*g*_st_). Thus, the *g*_liq_ was given as:

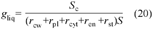

where *r*_cw_, *r*_pl_, *r*_cyt_, *r*_en_, *r*_st_ are the reciprocal term of *g*_cw_, *g*_st_, *g*_cyt_, *g*_en_, *g*_st_, respectively. Alternatively, according to Tholen *et al*.(2012b), the CO_2_ diffusion inside the cell in different ways: one for cell wall parts with chloroplast (*g*_cel,1_), and the other for inter-chloroplast areas (*g*_cel,2_), and the corresponding resistance was expressed as *r*_cel,1_ and *r*_cel,2_. Thus, the equation of *g*_liq_ was converted as:

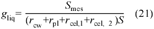

In addition, the conductance of liquid-phase diffusion pathway, either for cell wall (*g*_cw_), cytosol (*g*_cyt_), or stroma conductance (*g*_st_) is given as follows:

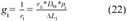

where *r*_f,i_ (dimensionless factor) accounts for the decrease of CO_2_ diffusion in the aqueous phase compared with free diffusion in water and was taken as 1.0 for cell wall and 0.3 for cytosol and stroma, as reported by Rondeau-Mouro *et al*.(2018) and Niinemets *et al*.(2003). Δ*L*_i_ (m) is the diffusion path length in the corresponding component of the diffusion pathway, and *p*_i_ (m^3^ m^−3^) is the effective porosity in the given part. The value of *p*_i_ was set to 1.0 for cytosol and stroma, and 0.29 for the cell wall. *D*_w_ is the CO_2_ diffusion coefficient in aqueous phase (1.79×10^−9^ m^-2^ s^-1^ at 25°C). Besides, conductance of plasma membrane (*g*_pl_) and chloroplast envelope (*g*_env_) was assumed as constant value of 0.0035 m s^-1^ (Evans *et al*., 1994; Tosens *et al*., 2012b).

### The quantitation of the anatomical limitation of the *g*_m_

The determinants of *g*_m_ were shared by gas-phase conductance (*l*_ias_) and liquid-phase conductance(*l*_i_). The proportion limited by *l*_ias_ was calculated as:

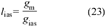

The share of *g*_m_ by the cellular-phase conductance (*l*_i_), which stands for the limitation of cell wall, the plasmalemma, and inside the cells was determined as:

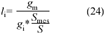

where *g*_i_ is the component diffusion conductance of the corresponding pathways. The fraction of exposed cell wall area lined with chloroplasts and fraction free of chloroplast were used to weighted limitations imposed by different cellular components (cytoplasm, chloroplast envelope, and stroma).

### Statistical analysis

Statistical analysis was conducted with SPSS 25.0 (SPSS Inc., Chicago, IL, USA). All data were subjected to a one-way analysis of variance (ANOVA), and the significant differences between treatments were compared using the least significant difference (*LSD*) at *P* < 0.05. Linear regression analyses were used to obtain the relationships among photosynthetic capacity and the main limiting factors, the key structural parameters and mesophyll conductance, and the values of mesophyll conductance estimated from different methods. Graphics and regression analyses were performed using Origin Pro 2021 software (Origin Lab Corporation, Northampton, MA, USA).

## Results

### Leaf morphology characteristics

Sole NH_4_^+^ supply significantly decreased leaf area (*A*_L_) and increased leaf thickness (*T*_L_) in comparison with sole NO_3_^−^ and mixed N supply (Table 1). Correspondingly, the leaf volume per area (*V*_A_) of sole NH_4_^+^ supply was almost half of the *V*_A_ of mixed N supply. The variation in leaf density (*D*_L_) was 1.5-fold with sole NH_4_^+^ supply having the densest leaves (0.36 g cm^-3^) and sole NO_3_^−^ supply the least dense (0.27 g cm^-3^). Notably, *M*_A_ seemed uninfluenced among the treatments. The leaf N concentration (*N*_L_) of sole NO_3_^−^ supply was approximately 11% lower compared with mixed N and sole NH_4_^+^ supply, yet the photosynthetic N use efficiency was dramatically upregulated by NO_3_^−^ treatments.

**Table 1.** Effect of different N forms on integral leaf variables

| Treatment | $M_A$ (mg cm<sup>-2</sup>) | $V_A$ (cm<sup>3</sup>) | $T_L$ (μm) | $D_L$ (g cm<sup>-3</sup>) | $A_L$ (cm<sup>2</sup>) | Chl (mg g<sup>-1</sup>) | $N_a$ (g m<sup>-2</sup>) | PNUE (μmol g<sup>-1</sup> s<sup>-1</sup>) |
| --- | --- | --- | --- | --- | --- | --- | --- | --- |
| A | 4.45±0.49a | 0.10±0.02b | 114.7±10.5a | 0.36±0.06a | 8.43±1.48b | 4.01±0.76a | 1.53±0.12a | 9.52±1.97c |
| AN | 4.76±1.46a | 0.20±0.02a | 106.5±9.6b | 0.40±0.12a | 13.81±1.83a | 4.18±0.42a | 1.43±0.21a | 13.16±1.70b |
| N | 4.01±0.71a | 0.19±0.03a | 108.9±9.9b | 0.27±0.04b | 13.32±0.93a | 4.36±0.60a | 1.18±0.11b | 16.87±1.80a |

$M_A$ , leaf mass per area; $V_A$ , leaf volume per area; $T_L$ , leaf thickness; $D_L$ , leaf density; $A_L$ , leaf area; Chl, total chlorophyll concentration per leaf; $N_a$ , leaf nitrogen content per area; PNUE, photosynthetic nitrogen use efficiency, $PNUE = A/N_a$ . A, AN, and N represent the sole $NH_4^+$ supply, mixed N supply, and sole $NO_3^-$ supply, respectively. Data are mean ± SD of 4 replications. Different letters indicate statistically significant differences ( $P < 0.05$ ).

### Leaf physiological characteristics

In comparison to sole NH_4_^+^ supply, net photosynthetic rate (*A*) was increased by both sole NO_3_^−^ and mixed N supply, by 39.2% and 26.6%, respectively. Whilst the intercellular CO_2_ concentration (*C*_i_) seemed indifferent among the treatments, chloroplast CO_2_ concentration (*C*_c_) was significantly down-regulated by sole NH_4_^+^ supply. For the stomatal conductance (*g*_s_) and mesophyll conductance (*g*_m_), sole NO_3_^−^ and mixed N supply had higher value than sole NH_4_^+^ supply and coincided with elevated electron transfer rate (*J*) and carboxylation efficiency (*CE*) (Table 2).

**Table 2.** CO_2_ transmission characteristics of *L. Japonica* affected by different N forms

| Treatment | $A$ ($\mu\text{mol}\cdot\text{m}^{-2}\cdot\text{s}^{-1}$) | $g_s$ ($\text{mol m}^{-2} \text{s}^{-1}$) | $C_i$ ($\mu\text{mol mol}^{-1}$) | $C_c$ ($\mu\text{mol mol}^{-1}$) | $C_i-C_c$ ($\mu\text{mol mol}^{-1}$) | $g_m$ ($\text{mol m}^{-2} \text{s}^{-1}$) | $J$ ($\mu\text{mol m}^{-2} \text{s}^{-1}$) | $CE$ |
| --- | --- | --- | --- | --- | --- | --- | --- | --- |
| A | 14.3±0.9b | 0.23±0.05b | 277.0±14.6a | 160.8±13.6b | 107.3±16.2a | 0.13±0.03b | 122.8±8.4b | 0.070±0.005b |
| AN | 18.1±2.5a | 0.35±0.10a | 275.1±11.8a | 188.8±8.0a | 86.3±7.9b | 0.21±0.04a | 138.3±13.4ab | 0.085±0.006a |
| N | 19.9±0.5a | 0.35±0.01a | 266.8±3.1a | 177.7±2.0a | 89.1±4.2b | 0.22±0.01a | 156.4±4.4a | 0.095±0.011a |

$A$ , net photosynthetic rate; $g_s$ , stomatal conductance; $C_i$ , intercellular CO<sub>2</sub> concentration; $C_c$ , chloroplast CO<sub>2</sub> concentration; $C_i-C_c$ , CO<sub>2</sub> drawdown from sub-stomatal cavities ( $C_i$ ) to chloroplasts ( $C_c$ ); $g_m$ , mesophyll conductance; $J$ , electron transfer rate; $CE$ , carboxylation efficiency. A, AN, and N represent the sole NH<sub>4</sub><sup>+</sup> supply, mixed N supply, and sole NO<sub>3</sub><sup>-</sup> supply, respectively. Data are mean ± SD of 5 replications for $A$ , $g_s$ , $C_i$ , $C_c$ , $C_i-C_c$ , $g_m$ , $J$ , and of 4 replications for $CE$ ; Different letters indicate statistically significant differences ( $P < 0.05$ ).

In order to ascertain the photosynthetic component that had the largest effect on net assimilation rate, we performed a correlation analysis, and the net photosynthesis rate (*A*) was positively correlated with stomatal conductance (*g*_s_), mesophyll conductance (*g*_m_), and electron transport rate (*J*) (Figure 2). The CO_2_ drawdown (*C*_i_-*C*_c_) from sub-stomatal cavities (*C*_i_) to chloroplasts (*C*_c_) ranged from 86.3 to 107.3 μmol mol^-1^ and was higher in leaves with decreased *g*_m_ under sole NH_4_^+^ supply.

### Leaf anatomical traits

Among the leaf ultrastructural characteristics estimated from TEM (Table 3; Figure 1), the mesophyll thickness (*T*_mes_), the volume fraction of intercellular air space (*f*_ias_), mesophyll surface area (*S*_mes_/*S*), and the chloroplast surface area (*S*_c_/*S*) exposed to intercellular airspace per unit leaf area were significantly down-regulated by sole NH_4_^+^ supply. Notably, there were marked differences in cell wall thickness (*T*_cw_) among the treatments, for sole NO_3_^−^ supply exhibited the thinnest cell wall (0.16 μm), while sole NH_4_^+^ supply had the thickness cell walls with the maximum value of 0.25 μm. For chloroplast characteristics, the length (*L*_chl_) and thickness (*T*_chl_) of the chloroplasts, and the distance of the chloroplast from the cell wall (*L*_cyt_) seemed unaffected by N forms supply, whereas NH_4_^+^ supply significantly elevated the distance between adjacent chloroplasts (*D*_chl-chl_). Besides, chloroplasts amount in per sponge cell and palisade cell showed more specific changes among the treatments. In the comparison with the other treatments, chloroplasts of both sponge and palisade cell were dramatically up-regulated in sole NO_3_^−^ supply (Supplementary Figure S1).

**Figure 1.**
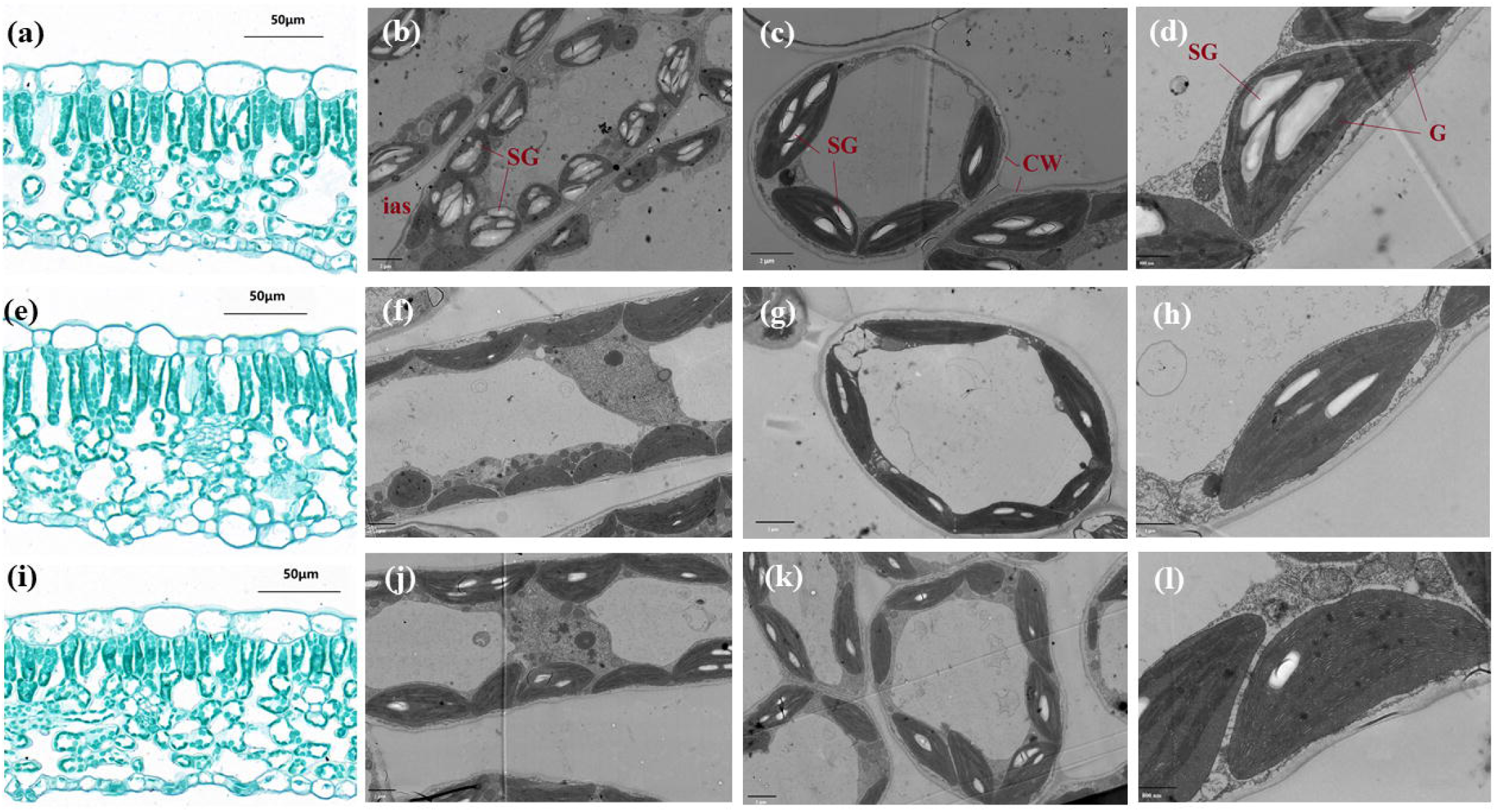
Light micrographs (a, e, i, scale bar = 50 μm) and transmission electron micrographs (b, f, j, c, g, k, scale bar = 2 μm; h, scale bar =1 μm; d, l, scale bar = 800 nm) of *L. Japonica* leaves affected by different forms of N, sole NH_4_^+^ supply (Fig. a-d), mixed N supply (Fig. e-h), and sole NO_3_^−^ supply (Fig. i-l). ias: intercellular air space; SG: starch granule; CW: cell wall; G: grana.

**Figure 2.**
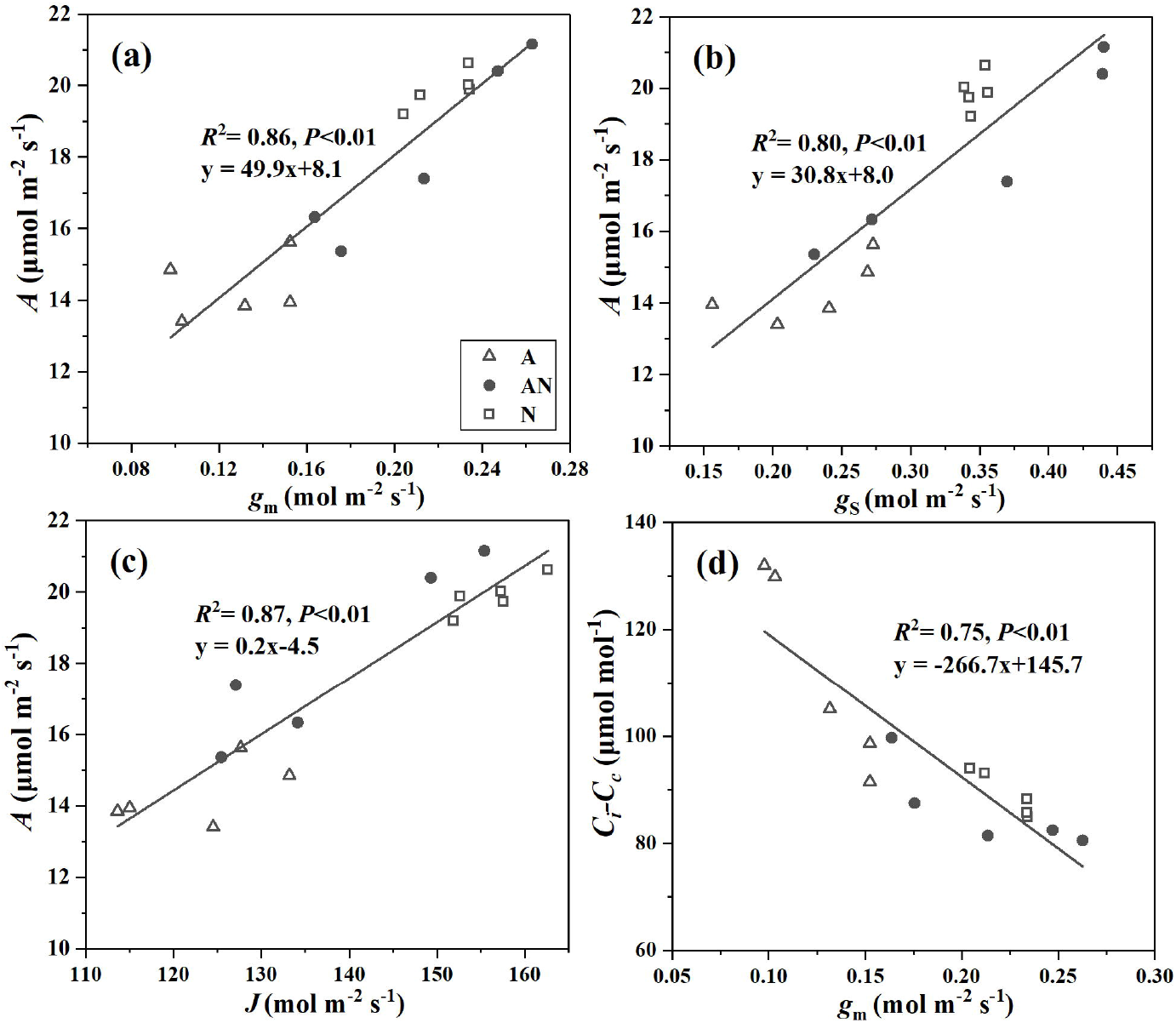
Correlation of the net photosynthetic rate(*A*) and mesophyll conductance (*g*_m_) **(a),** stomatal conductance (*g*_s_) **(b),** electron transport rate (*J*) **(c)**, and the relationship between CO_2_ drawdown (*C*_i_-*C*_c_) from sub-stomatal cavities (*C*_i_) to chloroplasts (*C*_c_) and *g*_m_ **(d)**. A, AN, and N represent the sole NH_4_^+^ supply, mixed N supply, and sole NO_3_^−^ supply, respectively, corresponding to the symbol of the open triangles, the closed circles, and the open square in the plot. The data were fitted by linear regressions (*P* < 0.01).

**Table 3.**
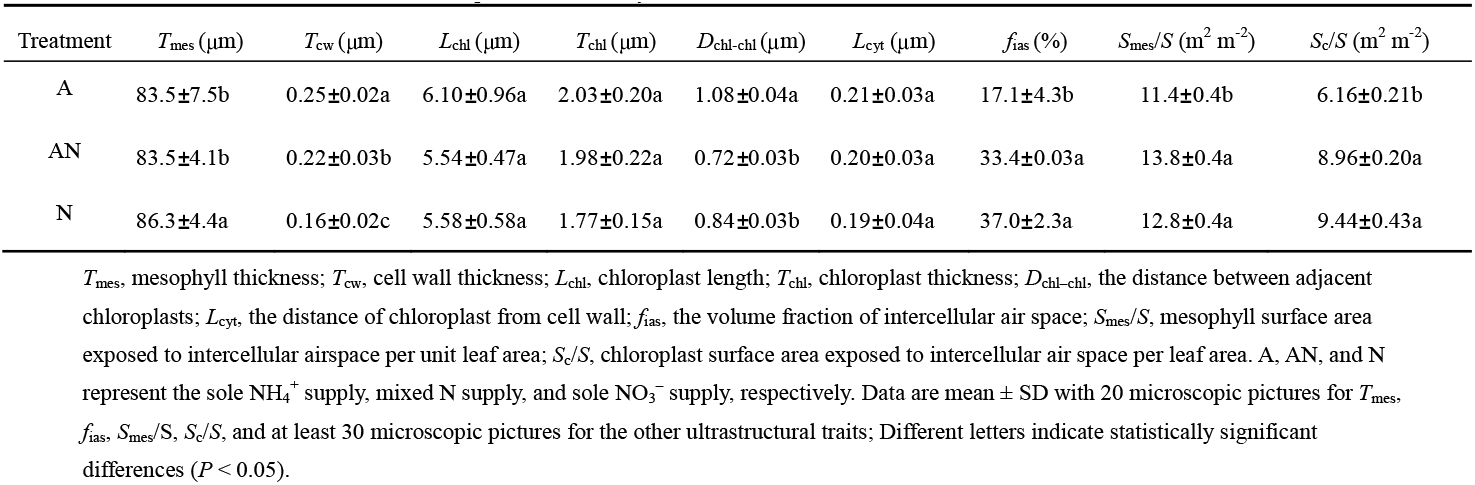
Leaf anatomical structure of *L. Japonica* affected by different N forms

| Treatment | $T_{\text{mes}}$ ($\mu\text{m}$) | $T_{\text{cw}}$ ($\mu\text{m}$) | $L_{\text{chl}}$ ($\mu\text{m}$) | $T_{\text{chl}}$ ($\mu\text{m}$) | $D_{\text{chl-chl}}$ ($\mu\text{m}$) | $L_{\text{cyt}}$ ($\mu\text{m}$) | $f_{\text{ias}}$ (%) | $S_{\text{mes}}/S$ ($\text{m}^2 \text{m}^{-2}$) | $S_{\text{c}}/S$ ($\text{m}^2 \text{m}^{-2}$) |
| --- | --- | --- | --- | --- | --- | --- | --- | --- | --- |
| A | 83.5±7.5b | 0.25±0.02a | 6.10±0.96a | 2.03±0.20a | 1.08±0.04a | 0.21±0.03a | 17.1±4.3b | 11.4±0.4b | 6.16±0.21b |
| AN | 83.5±4.1b | 0.22±0.03b | 5.54±0.47a | 1.98±0.22a | 0.72±0.03b | 0.20±0.03a | 33.4±0.03a | 13.8±0.4a | 8.96±0.20a |
| N | 86.3±4.4a | 0.16±0.02c | 5.58±0.58a | 1.77±0.15a | 0.84±0.03b | 0.19±0.04a | 37.0±2.3a | 12.8±0.4a | 9.44±0.43a |

$T_{\text{mes}}$ , mesophyll thickness; $T_{\text{cw}}$ , cell wall thickness; $L_{\text{chl}}$ , chloroplast length; $T_{\text{chl}}$ , chloroplast thickness; $D_{\text{chl-chl}}$ , the distance between adjacent chloroplasts; $L_{\text{cyt}}$ , the distance of chloroplast from cell wall; $f_{\text{ias}}$ , the volume fraction of intercellular air space; $S_{\text{mes}}/S$ , mesophyll surface area exposed to intercellular airspace per unit leaf area; $S_{\text{c}}/S$ , chloroplast surface area exposed to intercellular air space per leaf area. A, AN, and N represent the sole $\text{NH}_4^+$ supply, mixed N supply, and sole $\text{NO}_3^-$ supply, respectively. Data are mean ± SD with 20 microscopic pictures for $T_{\text{mes}}$ , $f_{\text{ias}}$ , $S_{\text{mes}}/S$ , $S_{\text{c}}/S$ , and at least 30 microscopic pictures for the other ultrastructural traits; Different letters indicate statistically significant differences ( $P < 0.05$ ).

Besides, *g*_m_ was not correlated with *T*_mes_, reflecting the circumstance that *T*_mes_ seemed more invariable. The positive and significant correlation between *g*_m_ and *S*_c_/*S, f*_ias_ were observed, combined with the negative relationship observed between *g*_m_ and *T*_cw_ suggested the importance of anatomical components to the intercellular CO_2_ diffusion.

### Estimation of *g*_m_ with different methods

In the present study, the value of *g*_m_ was measured according to the gas exchange and chlorophyll fluorescence (Harley *et al*., 1992), and compared with *g*_m_ modelled by *A/C*_i_ response curves (Bernacchi *et al*., 2002) and anatomical characteristics (Tomás *et al*., 2013). The correlation analysis indicated that the estimated *g*_m_ from different methods was mainly positively correlated (Supplementary Figure S3). However, the slope was different from unity, and the Harley *et al*.-based and Tomás *et al*.-based estimates of *g*_m_ were strongly correlated with a *R*^2^ of 0.76 (*P* < 0.01).

### Limiting components analyses of *A*

The relative limitations of stomatal conductance(*l*_s_), mesophyll diffusion (*l*_m_), and biochemistry capacity (*l*_b_) on photosynthesis are shown in Figure 3(a). In the NO_3_^−^ treatment and mixed N treatment, the percentages of the three limiting components are relatively close, and *l*_b_ (36.4%and 39.5%, respectively) accounts for the most important relative limitation of photosynthesis. While for sole NH_4_^+^ supply, the relative limitation of *l*_s_, *l*_m_, and *l*_b_ significantly varied, and the *l*_m_ (44.2%) was considered more important of photosynthesis, followed by *l*_s_ (30.8%) and *l*_b_ (25.0%). Meanwhile, according to quantitative limitations analysis of *A*, mesophyll conductance limitation (*MC*_L_) was the highest constrain of *A* under sole NH_4_^+^ supply, accounting for the 24.39 % of *A* decline compared with the NO_3_^−^ treatment (Figure 3).

**Figure 3.**
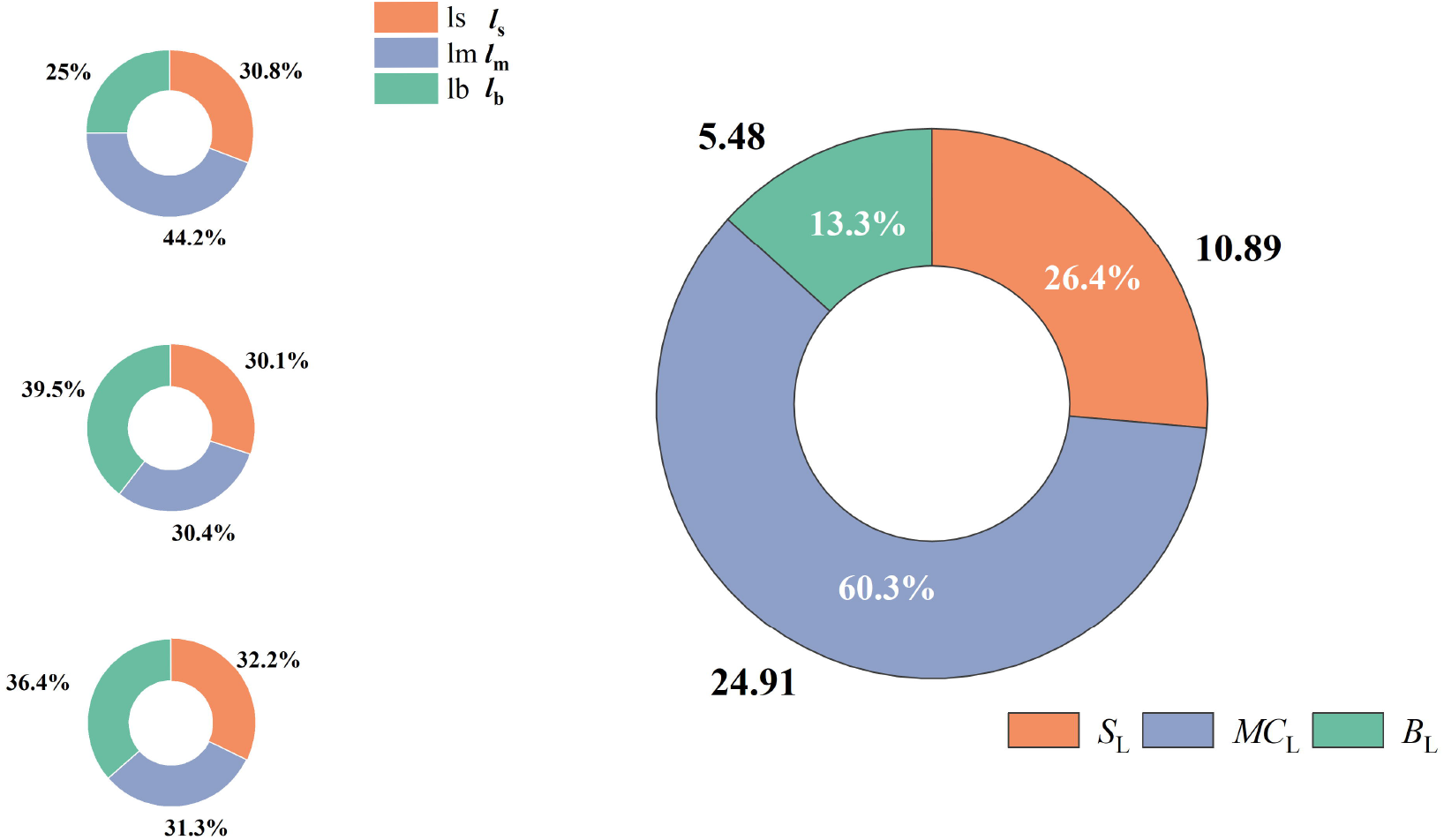
Relative limitation of stomatal conductance (*l*_s_), mesophyll diffusion (*l*_m_), and biochemical capacity (*l*_b_) of *L. Japonica* under sole NH_4_^+^ supply **(a)**, mixed N supply **(b),** and sole NO_3_^−^ supply treatment **(c)**, the limitation of *l*_s_, *l*_m_, and *l*_b_ together add up to 100% at each treatment occasions; Quantitative limitation analyses of stomatal limitation (*S*_L_), mesophyll conductance limitation (*MC*_L_), and biochemical limitation (*B*_L_) of *L. Japonica* photosynthesis under sole NH_4_^+^ supply **(d)**, the data outside the circles represent the absolute quantitative limitations and the data inside the circles represent the share of the total limitations by *S*_L_, *MC*_L_, and *B*_L_.

### Key structural factors regulating *A* through *g*_m_

For the different components of the whole CO_2_ diffusion pathway, restrictions in liquid-phase CO_2_ conductance were the dominant limiting factors on *g*_m_, accounting for approximately 90% of the total limitation (Figure 4b). Among the different components of the liquid-phase, the stroma limitation was the main limiting factor in the liquid-phase, whereas the stroma limitation seemed unaffected by the treatments. The liquid-phase limiting component of *g*_m_ is also associated with the cytoplasm, with significantly upregulated by sole NH_4_^+^ supply. In comparison with the sole NH_4_^+^ supply, plasma membrane and chloroplast envelope limitation proposed a slightly higher percentage in sole NO_3_^−^ supply, with 24.3% and 24.3%, respectively. For the absolute value of the limitations, the cell walls appeared to have a more specific change among the treatments that limited the internal diffusion of CO_2_ varied from 303.69 to 480.37 s m^-1^ (Figure 4a). Besides, the variation in cytoplasm resistance was 1.7-fold, with sole NO_3_^−^ treatment having the lowest resistance (450.13 s m^-1^) and sole NH_4_^+^ supply having the highest (664.23 s m^-1^).

**Figure 4.**
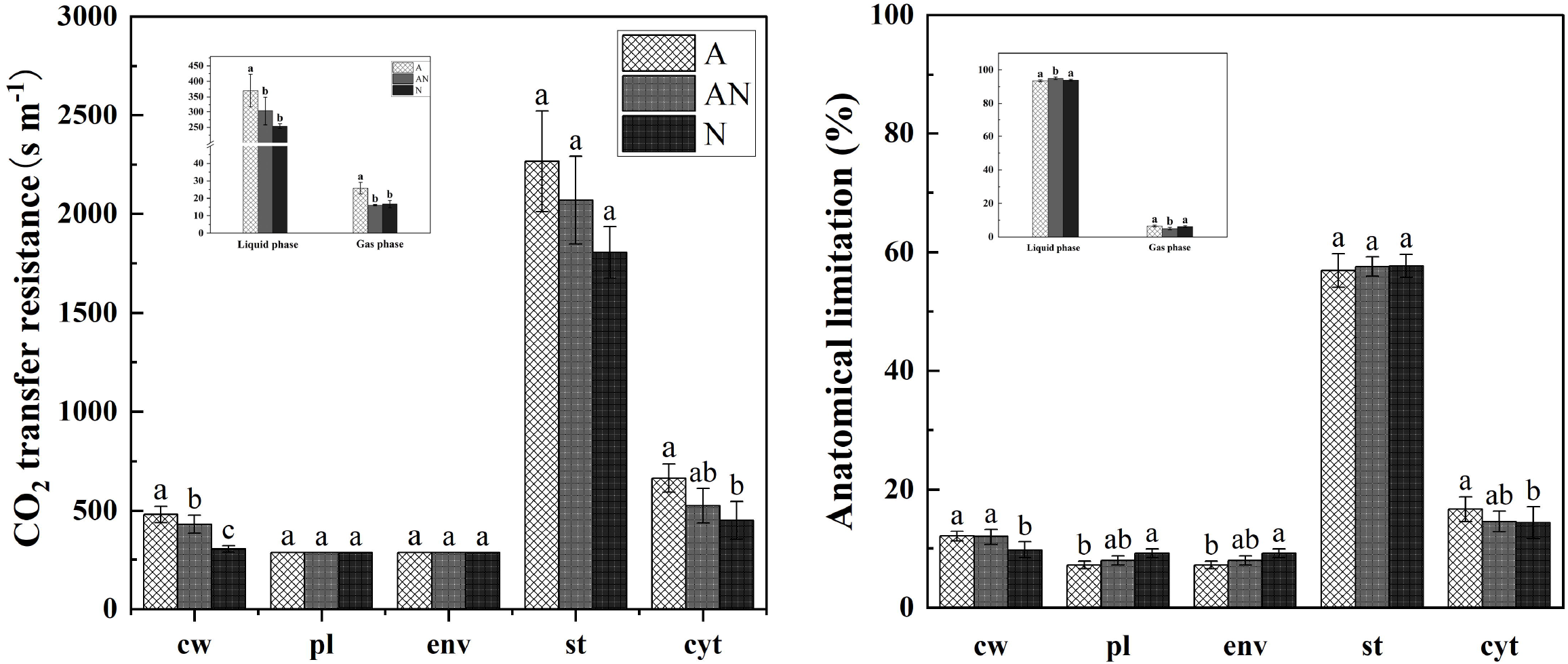
Anatomical limitations of mesophyll conductance (*g*_m_) **(a)** and the share of the overall *g*_m_ limitation **(b)** by cell wall (cw), plasma membrane (pl), chloroplast envelope (env), stroma (st), and cytoplasm (cyt) in leaves of *L. Japonica* under different forms of N. A, AN, and N represent the sole NH_4_^+^ supply, mixed N supply, and sole NO_3_^−^ supply, respectively. The inset figures showed the anatomical limitations of *g*_m_ and the share of the overall *g*_m_ limitation by gas-phase and liquid-phase. Different letters indicate statistically significant differences (*P* < 0.05).

## Discussion

### *g*_m_ dominated the decrease of *A* in sole NH_4_^+^ supply

In the present study, inorganic N sources significantly decreased the photosynthetic rate (*A*) of *Lonicera Japonica*. Leaf in sole NH_4_^+^ treatment was observed a dramatically downregulated *A*, in parallel with a larger dropdown of CO_2_ from the sub-stomatal cavities (*C*_i_) to the sites of carboxylation inside the chloroplasts (*C*_c_). Among the limiting factors, mesophyll conductance limitation (*M*C_L_) controlled 60.3 % of *A* decline, followed by stomatal conductance (*S*_L_, 26.4%) and biochemistry limitation (*B*_L_, 13.3%). These results suggested the mesophyll diffusion resistance to CO_2_ is a key limiting factor to *A*. Early studies suggested that *g*_m_ was driven by integrated leaf characteristics, notably and negatively correlated with leaf mass per area (*M*_A_) (Flexas *et al*., 2008; Han, 2011; Niinemets *et al*., 2009a; Onoda *et al*., 2017). Nevertheless, the *M*_A_ did not show any significant difference among treatments in this study, despite some declining tendency of sole NO_3_^−^ treatment. Whist the unchanged *M*_A_, the present data showed that the NH_4_^+^-fed leaves were manifested by higher leaf thickness (*T*_L_) and density (*D*_L_), and a smaller leaf area (*A*_L_) and leaf volume (*V*_A_) (Table 1). The leaf traits are generally observed in NH_4_^+^-fed plants compared to those supplied with NO_3_^−^ (Guo *et al*., 2007), suggesting the high leaf structure plasticity and varied leaf carbon-expensive structure in relation to environmental conditions.

As there is ample agreement that the use of integrated traits such as *M*_A_ as proxies of *g*_m_ might not be valid in all cases, the intercellular anatomical traits that limit CO_2_ effective diffusion length and area, in particularly the cell wall thickness (*T*_cw_) are especially crucial (Evans *et al*., 2009; Momayyezi and Guy, 2017; Tosens *et al*., 2012b). In the present study, a one-dimensional within-leaf gas diffusion model considering all of the leaf anatomical limitations was applied as previously modified by Tomas *et al*. (2013). The *g*_m_ modelled from anatomical traits was tightly correlated with measured *g*_m_ (Supplementary Figure S3), which, to some extent, verified the view that the variation of *g*_m_ is related to the intercellular structures involved in CO_2_ diffusion. Nevertheless, the partial conductance components are generally assumed to be composed of a single medium in the estimation of *g*_m_, which might be unrealistic (Gago *et al*., 2020). In addition, the carbonic anhydrases, as well as aquaporins were not considered in the *g*_m_-modelled (Flexas *et al*., 2012). Consequently, the correlation between *g*_m_-gas exchanges and *g*_m_-modelled frequently deviates from the 1:1 relationship.

Mesophyll conductance is a composite conductance of an intercellular gas-phase (*g*_ias_) and a liquid-phase (*g*_liq_). The *g*_ias_ mainly depends on the intercellular air space and mesophyll thickness, while the *g*_liq_ was affected by apoplast and cellular components of the CO_2_ pathway, which can be scaled by the chloroplast surface area exposed to intercellular air space per unite leaf area (*S*_c_/*S*) (Flexas *et al*., 2012). It was observed that the limitations were mainly represented by liquid-phase, and the gas-phase accounts for only approximately 10% (Figure 4b), analogous to the value obtained in rape (Lu *et al*., 2016). Despite the low proportion, a significant discrepancy of *g*_ias_ among the treatments appeared, in that gas-phase limitation was upregulated by sole NH_4_^+^ supply. This result was somewhat inconsistent with Liu *et al* (2021) and Gao *et al* (2020), as NO_3_^−^ caused higher limitation on *g*_ias_. This could be explained by the variation of the environment light intensity, concentrations of N addition, experiment period as well as the plant species. Integrated all the determinants into account, there was a positive correlation between *g*_m_ and the volume fraction of intercellular air space (*f*_ias_) (*R*^2^ = 0.71, *P* < 0.01), while a weak relationship with the mesophyll thickness (Figure 5), suggesting that *f*_ias_ was more variable in regulating *g*_m_. Additionally, *f*_ias_ under sole NO_3_^−^ supply was approximately 2-folds higher than that of sole NH_4_^+^ supply, similar to the result of Liu *et al* (2021). The flexible *f*_ias_ were general in plants, which may be ascribed to acclimation-related changes, e.g., a larger *f*_ias_ was usually observed in more favorable growth conditions (Binks *et al*., 2016; Muller *et al*., 2009).

**Figure 5.**
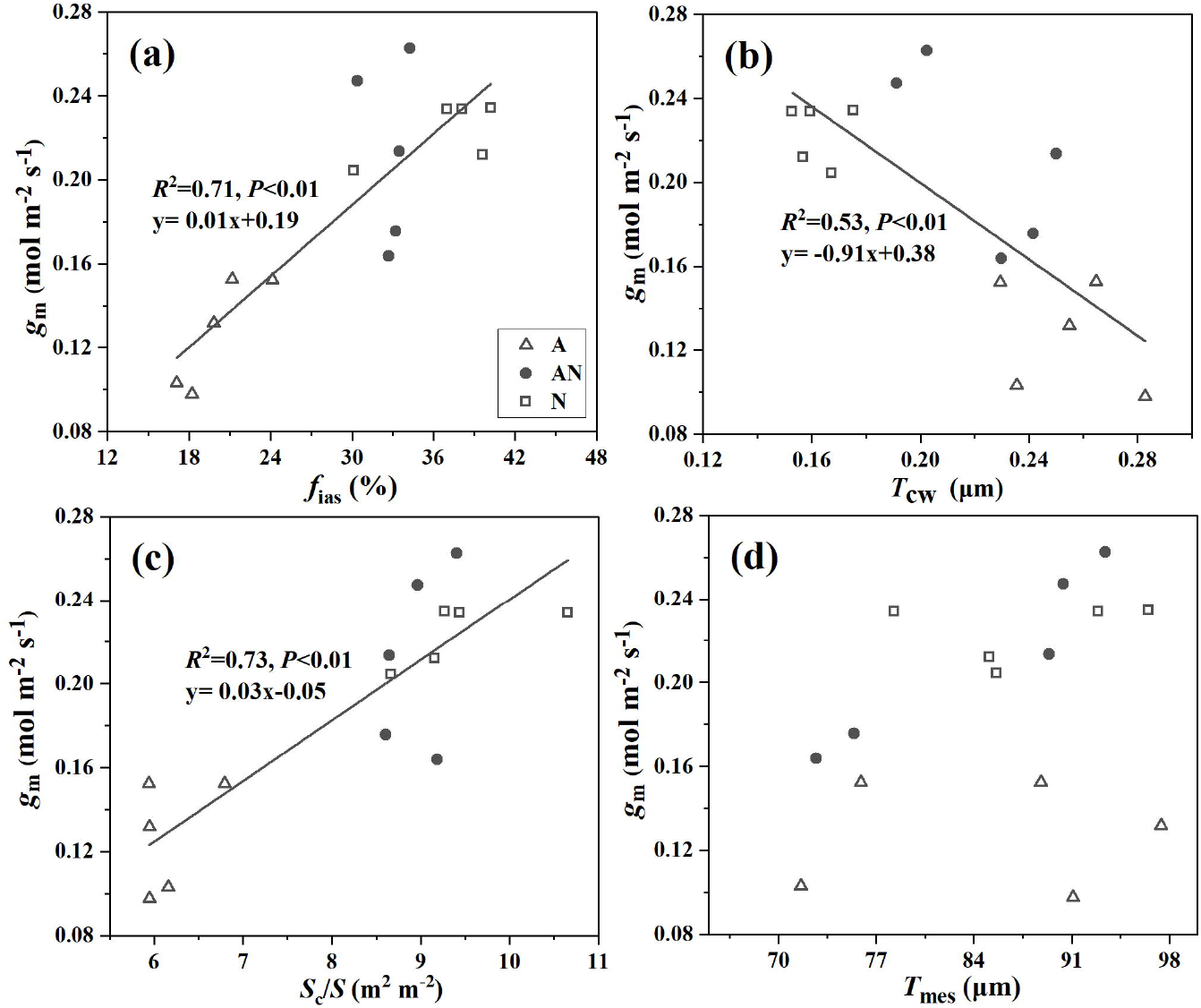
Correlations of mesophyll conductance (*g*_m_) with the intercellular air space (*f*_ias_) **(a)**, the volume fraction of cell wall thickness (*T*_cw_) **(b),** the chloroplast surface area exposed to intercellular air space per leaf area (*S*_c_/*S*) **(c)**, and mesophyll thickness (*T*_mes_) **(d)**. A, AN, and N represent the sole NH_4_^+^ supply, mixed N supply, and sole NO_3_^−^ supply, respectively, corresponding to the symbol of the open triangles, the closed circles, and the open square in the plot. The data of each group was fitted by linear regressions (*P* < 0.01).

### Structural trade-off in driving the share of *A* limitations

As suggested in various studies, *S*_c_/*S* is more influential in determining *g*_m_ across species (Loucos *et al*., 2017; Ren *et al*., 2019; Terashima *et al*., 2011; Veromann-Jürgenson *et al*., 2020; Veromann-Jurgenson *et al*., 2017). Here, the down-regulation of *S*_c_/*S* by sole NH_4_^+^ supply was speculated to be attributed to the fewer chloroplast numbers and the lower mesophyll surface area exposed to intercellular airspace per leaf area (*S*_mes_/*S*) (Table 3), similar to the observation of (Carriqui *et al*., 2015); while the length and thickness of chloroplasts were unchanged among the treatments. Despite the dropdown of *r*_st_, a fact that cannot be ignored is the relative control of chloroplast stroma on *g*_liq_ was less affected by N forms, with no significant change among the treatments.

There rises a crucial concern that whether *g*_m_ was necessarily controlled by the component owning the largest proportion, because the contribution of components that limit CO_2_ diffusion may vary among species. Here, the discrepancy in liquid-phase resistance was suggested to be related to the cytoplasm and cell wall. For cytoplasm, it accounts for approximately 15% of liquid resistance in the present study, corresponds well with previous reports (Lu *et al*., 2016; Tosens *et al*., 2012a; Tosens *et al*., 2012b). Commonly, the chloroplasts are arranged against the cell periphery to absorb light and CO_2_ (Sage *et al*., 2009), thus the distance of chloroplast from cell wall (*L*_cyt_) could be responsible for most of the cytoplasm resistance, as reported by (Sharkey *et al*., 1991). However, *L*_cyt_ was the same among the treatments (Table 3) in this study, and therefore, the upregulated *r*_cyt_ in NH_4_^+^ treatment may be explained by the increase of distance between adjacent chloroplasts (*D*_chl-chl_), by up to 50% compared with mixed N treatment.

Previous studies have highlighted the importance of cell wall in determining *g*_m_ among species, as the cell wall could account for 50% of *g*_m_ by restricting CO_2_ diffusion (Evans *et al*., 2009; Terashima *et al*., 2006). Although the limitation is often neglected, cell wall limitation sometimes is greater than *S*_c_/*S*, and together these constitute the primary anatomical factors for setting the maximum *g*_m_ (Carriquí *et al*.,2019; Carriqui *et al*., 2019). Here, the cell wall thickness (*T*_cw_) varied over a range of 0.16 to 0.25 μm, and *T*_cw_ of sole NO_3_^−^ treatment decreased by 27% to 36% in the comparison with mixed N and sole NH_4_^+^ supply, resulting in a dramatic difference among the treatments. Głazowska *et al*. (2019) had reported that the distinct cell wall remodeling was mediated by inorganic N supply. NH_4_^+^-mediated cell wall thicken, as observed in early studies may be ascribed to changes in the contents of polysaccharide, ion, lignin, or cellulose of cell walls (Ellsworth *et al*., 2018; Podgórska *et al*., 2017). Meanwhile, as previously mentioned, a more rigid cell wall structure could explain the reduction in *A*_L_ in ammonium supply, by limiting cell expansion. Besides, the *T*_cw_ and *g*_m_ were observed strongly and positively correlation in this study (Figure 5 (b), *R*^2^ = 0.53, *P* < 0.01), the reduced *g*_m_ and thicker cell wall were reported to be an adaption of plant species to dry or nutrition poor environment (Niinemets *et al*., 2009a; Niinemets *et al*., 2009b).

Recent evidence points to an effect of cell wall composition on photosynthesis, possibly due to a trade-off of N allocation between chloroplasts and the cell wall in plants (Kuusk *et al*., 2018). In angiosperms, particular in C3 plants, chloroplast N distribution accounts for almost 75 %, while the cell wall for 10% (Li *et al*., 2017; Onoda *et al*., 2017; Wang *et al*., 2015); the leaf N that is not allocated to photosynthetic apparatus is generally used structurally in cell walls (Feng *et al*., 2009). In this study, the reduced chloroplasts numbers and the thicker *T*_cw_ by sole NH_4_^+^ supply were speculated to be related to the down-regulated photosynthetic N allocation (Figure S1). Takashima *et al* (2004) had reported that higher N allocation to the cell wall could lead to decreased PNUE. The discrepancy of N partitioning among N sources may indicate a trade-off in the leaf photosynthetic capacity and the persistence, while the mechanism underlying this result need further research.

## Conclusion

The present study showed that N sources significantly affected the leaf morphology and photosynthetic rate (*A*) of *L. Japonica* and suggested the mesophyll diffusion resistance accounts for the most limiting of *A*, with more than 50%. Sole NH_4_^+^-fed leaves were characterized by smaller leaf area, higher leaf thickness, and larger leaf density. Variations of mesophyll conductance (*g*_m_) under different N sources were ascribed to the leaf anatomy changes, notably the internal air space (*f*_ias_), exposed surface area of chloroplasts per unit leaf area (*S*_c_/*S*) and cell wall thickness (*T*_cw_). Ammonium treatment reduced the *f_ias_* and chloroplast numbers, resulting in an increased intercellular length and inter-chloroplast length, and finally inhibition of *g*_m_ and *A* (Figure 6).

**Figure 6.**
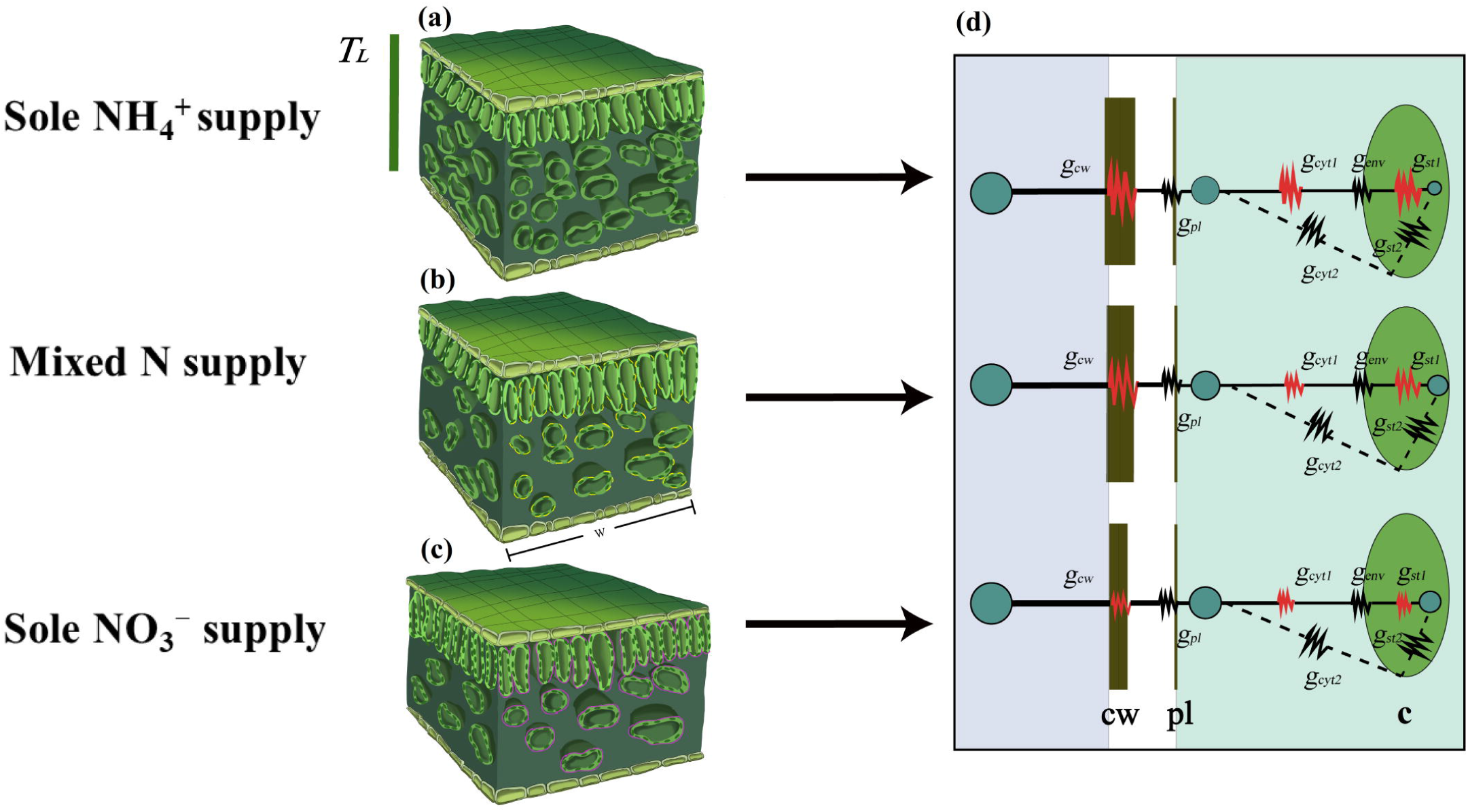
A schematic model of *L. Japonica* leaf anatomical traits and CO_2_ diffusion pathway under different forms of N supply. Leaf ultrastructure model in sole NH_4_^+^ supply **(a)**, mixed N supply **(b),** and sole NO_3_^−^ supply **(c),** respectively. CO_2_ diffusion pathways inside the mesophyll cells in sole NH_4_^+^ supply, mixed N supply, and sole NO_3_^−^ supply, respectively **(d)**. The chloroplast surface area exposed to intercellular air space per leaf area (*S*_c_/*S*) was outlined with yellow lines in (b), and the mesophyll surface area exposed to intercellular airspace per unit leaf area (*S*_mes_/*S*) was marked with purple lines in (c). The loose arrangement of mesophyll cells as affected by sole NO_3_^−^ supply increased the intercellular airspace and consequently up-regulated the *S*_c_/*S* and *S*_mes_/*S* **(b, c)**. Increased chloroplast density resulted in reduced distance between two adjacent chloroplasts (*D*_chl-chl_) in sole NO_3_^−^-fed leaves **(c)**. The red folded lines and black folded lines represent the strength of CO_2_ diffusion resistance into the cell from cell wall (*g*_cw_), plasma (*g*_pl_), cytoplasm (*g*_cyt_), envelope (*g*_env_) and stroma (*g*_st_), while the red folded lines indicated the values of this component differed between treatments and the black folded lines means no difference between treatments or not been measured in the study **(d)**. The blue dot represents the CO_2_ concentration **(d)**. *T*_L_, leaf thickness; W, the width of the leaf section; cw, cell wall; pl, plasm; c, chloroplast.

## Acknowledgments

This work was financially supported by the National Natural Science Foundation of China (32072673), the Fundamental Research Funds for the Central Universities (KYGD202007), the Young Elite Scientists Sponsorship Program by CAST (2018QNRC001), and the Innovative Research Team Development Plan of the Ministry of Education of China (IRT_17R56).

## Author contributions

Shiwei Guo and Yiwen Cao conceived the idea and designed the experiment; Yiwen Cao, Yonghui Pan, and Tianheng Liu completed the experiment; Yiwen Cao analyzed the data and wrote the manuscript; Shiwei Guo, Min Wang, and Yonghui Pan helped in manuscript revising; Shiwei Guo and Min Wang provided funding support. All the authors contributed critically to the drafts and gave final approval for publication.

